# The green tea catechin EGCG provides proof-of-concept for a pan-coronavirus entry inhibitor

**DOI:** 10.1101/2021.06.21.449320

**Authors:** Emmanuelle V. LeBlanc, Che C. Colpitts

## Abstract

The COVID-19 pandemic caused by the severe acute respiratory syndrome coronavirus-2 (SARS-CoV-2) has emphasized the serious threat to human health posed by emerging coronaviruses. Effective antiviral countermeasures are urgently needed as vaccine development and deployment against an emerging coronavirus takes time, even in the best-case scenario. The green tea catechin, epigallocatechin gallate (EGCG), has broad spectrum antiviral activity. We demonstrate here that EGCG prevents murine and human coronavirus infection and blocks the entry of lentiviral particles pseudotyped with spike proteins from highly pathogenic coronaviruses, as well as a bat coronavirus poised to emerge into humans. We show that EGCG treatment reduces coronavirus attachment to target cell surfaces. Our results demonstrate the potential for the development of pan-coronavirus attachment inhibitors to protect against current and future emerging coronaviruses.

## 1. Introduction

The novel severe acute respiratory syndrome coronavirus-2 (SARS-CoV-2) has infected hundreds of millions of people and has resulted in millions of deaths around the world. The COVID-19 pandemic, caused by SARS-CoV-2, underscores the severe health threat posed by emerging coronaviruses (CoVs), and makes evident the paucity of antivirals with extended spectrums of activity. SARS-CoV-2 is the third highly pathogenic CoV to emerge in the 21^st^ century. The abundance and diversity of SARS-related CoVs in bats points to the likelihood of future spillover into human populations (Li et al., 2005). In addition, seasonally circulating common cold CoVs continually burden human health (Walsh et al., 2013), and cause severe disease in elderly populations (Hand et al., 2018; Patrick et al., 2006), without effective preventative or therapeutic strategies. In order to develop inhibitors that are effective against such a broad range of endemic, pandemic, and emerging CoVs, a conserved step of viral infection needs to be targeted.

Most human viruses, including CoVs, initiate primary attachment to cell surfaces through low-affinity but high-avidity interactions with glycans. Heparan sulfate proteoglycans are necessary attachment receptors for the human alphacoronavirus NL63 (HCoV-NL63) (Milewska et al., 2014), as well as many other human viruses, such as herpes simplex virus (HSV) (Lycke et al., 1991; WuDunn and Spear, 1989), hepatitis C virus (HCV) (Germi et al., 2002; Morikawa et al., 2007) and human immunodeficiency virus (HIV) (Patel et al., 1993). Recently, heparan sulfate has also been identified as a necessary co-factor for SARS-CoV-2 infection (Clausen et al., 2020; Zhang et al., 2020). Alternatively, other viruses, including influenza A virus (IAV) and betacoronaviruses HCoV-OC43 and HCoV-HKU1 require sialic acid-containing glycans for attachment and entry (Huang et al., 2015; Krempl et al., 1995; Weis et al., 1988). Regardless of the glycan subtype, glycan binding is critical to the primary attachment of many different viruses, facilitating their specific high-affinity interaction with less abundant protein receptors, such as ACE2 for HCoV-NL63, SARS-CoV-1 and SARS-CoV-2 (Hoffmann et al., 2020; Hofmann et al., 2005). For some viruses, like IAV, glycans serve as the only known entry receptor.

Blocking viral attachment to cellular glycans is a demonstrated approach to inhibit diverse viral infections. Epigallocatechin gallate (EGCG) is the most bioactive polyphenolic compound from green tea with antiviral activity against diverse DNA and RNA viruses, including HSV-1, HCV, and IAV (Xu et al., 2017). The broad antiviral spectrum of EGCG has been attributed to its ability to inhibit the primary attachment of unrelated viruses by competing with both heparan sulfate and sialic acid for virion binding (Colpitts and Schang, 2014). However, whether EGCG inhibits coronavirus infection by similar mechanisms remains unknown.

Here, we show that EGCG inhibits infectivity of murine, bat, and human CoVs. EGCG blocks cell surface binding of multiple CoVs, suggesting that it inhibits a conserved step in CoV attachment, such as initial binding to glycans. These findings demonstrate that blocking primary attachment is a potential antiviral strategy to prevent infection by diverse CoVs.

## 2. Material and Methods

### 2.1. Compounds

Epigallocatechin-3-gallate (EGCG, 95% pure) was purchased from Thermo Fisher Scientific (AC449010100). Heparin was obtained from Carbosynth Ltd (YH09354).

### 2.2. Plasmids

Lentiviral pseudoparticles were produced using plasmids encoding HIV-1 gag/pol (BEI Resources NR-52517), tat (BEI Resources NR-52518), rev (BEI Resources NR-52519), a luciferase-encoding lentiviral genome (BEI Resources NR-52516) and spike plasmids. The SARS-CoV-2 spike expression plasmid was obtained from Dr. Raffaele De Francesco (Addgene 155297). The MERS-CoV spike expression plasmid was kindly provided by Dr. Stefan Pöhlmann (Göttingen, Germany). The SARS-CoV-1 (Tor2 strain, GenBank accession no. **<u>NC_004718.3</u>**) and WIV1-CoV spike (GenBank accession no. **<u>KC881007.1</u>**) expression plasmids were synthesized by Genscript and codon-optimized for human expression. The lentiviral vector expressing human ACE2 (Dr. Sonja Best, Addgene 154981) was used with the lentiviral packaging plasmid psPAX2 (Dr. Didier Trono, Addgene 12260) and pCMV-VSV-G (Dr. Bob Weinberg, Addgene 8454).

### 2.3. Cells and viruses

HEK293T/17 (ATCC CRL-11268), L929 (ATCC CCL-1), Huh7 (JCRB0403), HCT-8 (ATCC CCL-244) and 17CL1 (BEI Resources NR-53719) cells were cultured in Dulbecco’s minimal essential medium (DMEM) with 10% FBS, 50 U/mL penicillin, and 50 μg/mL streptomycin at 37°C in 5% CO_2_. A549 (BEI Resources NR-52268) cells were cultured in Hams F-12K (Kaighn’s) medium with 10% FBS, 50 U/mL penicillin, and 50 μg/mL streptomycin at 37°C in 5% CO_2_. Calu-3 (ATCC HTB-55) cells were cultured in minimal essential medium (MEM) with 10% FBS, 1 mM sodium pyruvate, 1X non-essential amino acid solution, 50 U/mL penicillin, and 50 μg/mL streptomycin at 37°C in 5% CO_2_.

A549-ACE2 cells were generated by lentiviral transduction and selected with 10 μg/mL blasticidin. The bulk population was single-cell cloned by limiting dilution, and ACE2 expression of clonal populations was determined by Western blotting using a rabbit monoclonal ACE2 antibody (Invitrogen MA5-32307), mouse monoclonal actin (Abcam ab8226) or tubulin (Abcam ab7291) antibodies, and a LICOR Odyssey CLx with appropriate secondary antibodies.

Human coronavirus 229E (HCoV-229E) and HCoV-OC43 were obtained from BEI Resources (NR-52726 and NR-52725). For HCoV-229E, Huh7 cells were infected with a multiplicity of infection (MOI) of 0.01 plaque-forming units (pfu)/cell and incubated at 33°C in 5% CO_2_ until full cytopathic effects (CPE) were observed (∼5 days post-infection). The supernatants were collected, and cellular debris was pelleted by centrifugation at 1,000 × g for 10 min. The resulting supernatant was then aliquoted, and the viral stocks were stored at −80°C. For HCoV-OC43, HCT-8 cells were infected (MOI of 0.01) and incubated at 33°C in 5% CO_2_ until full CPE were observed (∼5 days post-infection). Similarly, supernatants were collected, cellular debris was pelleted by centrifugation at 1,000 × g for 10 min and the resulting supernatant was then aliquoted and stored at −80°C.

Green fluorescent protein-expressing murine hepatitis virus (MHV A59-GFP) was kindly provided by Dr. Volker Thiel (Institute of Virology and Immunology, Bern, Switzerland) (Coley et al., 2005). VSV-SARS-CoV-2 S was a gift from Dr. Sean Whelan (Washington University School of Medicine in St. Louis, USA) (Case et al., 2020). MHV A59-GFP and VSV-SARS-CoV-2 were propagated as described (Coley et al., 2005; Case et al., 2020).

### 2.4. Infectivity assays

Huh7 cells (1.6 × 10^6^ cells/well in 6-well plates) were infected with approximately 100 PFU of HCoV-229E pre-exposed for 10 min at 37°C to EGCG or dimethyl sulfoxide (DMSO) vehicle in DMEM. Inocula were removed 2 h later, and monolayers were overlaid with 1.2% carboxymethylcellulose (Fisher Scientific AAA1810536) in DMEM containing 2% FBS. Infected cells were incubated at 33°C in 5% CO_2_ until 4 days post-infection, when they were fixed with 10% formalin and stained with 1% (wt/vol) crystal violet–10% (vol/vol) ethanol in H_2_O for counting plaques.

Huh7 and A549 cells (4 × 10^5^ cells/well in 24-well plates) were infected with approximately 200 focus-forming units (FFU) of HCoV-OC43, pre-exposed to EGCG or DMSO vehicle for 10 min at 37°C in DMEM. Inocula were removed 2 h later, and monolayers were overlaid with 1.2% microcrystalline colloidal cellulose (Sigma-Aldrich 435244) in DMEM containing 2% FBS. Infected cells were incubated at 33°C in 5% CO_2_ for 3 days, then fixed and processed for immunofluorescence to detect OC43 nucleoprotein. Cells were incubated with primary mouse IgG anti-coronavirus group antibody MAB9013 (Millipore Sigma; diluted 1:500) for 1 h at room temperature, followed by addition of secondary Alexa Fluor 488 anti-mouse IgG Fab 2 antibody (Cell Signaling Technology 4408S) for 1 h at room temperature. Foci were visualized and counted under a fluorescence microscope (Nikon Eclipse Ts2).

MHV A59-GFP virions pre-exposed to EGCG or the DMSO vehicle control for 10 min at 37°C in DMEM were used to infect L929 cell monolayers. Inocula were removed 1 h later, and monolayers were overlaid with 1.2% carboxymethylcellulose in DMEM containing 2% FBS. Infected cells were incubated at 33°C in 5% CO_2_ for 24 h, then fixed prior to counting of GFP-positive cells under the fluorescence microscope.

VSV-SARS-CoV-2 S virions pre-exposed to EGCG or the DMSO vehicle control for 10 min at 37°C in DMEM were used to infect Huh7, A549-ACE2 or Calu-3 cell monolayers in duplicate wells of a 96-well plate. Infected cells were incubated at 33°C in 5% CO_2_ for 24 h, then fixed with 10% formalin. GFP signals were captured by Nikon Eclipse Ts2, processed using the Python Imaging Library to find the mean grey value, and plotted as percentage of inhibition.

In all experiments, half maximal inhibitory concentrations (IC_50_) were calculated by nonlinear regression analysis using GraphPad Prism (version 9.0; GraphPad Software, Inc.).

### 2.5. Pseudoparticle entry assays

Lentiviral pseudoparticles were generated in HEK293T/17 cells by co-transfection using Lipofectamine 2000 (Invitrogen 11668-019) (Crawford et al., 2020). The CoV spike expression plasmids were co-transfected with a luciferase-encoding lentiviral genome, and plasmids encoding HIV-1 gag/pol, tat, and rev (Crawford et al., 2020). Cell culture supernatants containing coronavirus spike pseudotyped lentiviral particles were collected at 48 and 72 hours post-transfection, pooled and filtered through a 0.45 μm filter, and stored at −80°C.

Pseudoparticles were incubated for 10 min at 37°C with serially diluted EGCG and used to infect Huh7 or A549-ACE2 cells in triplicate in 96-well plates. Infectivity was determined after 3 days by luminescence following addition of BrightGlo reagent (Promega PR-E2620). IC_50_ were calculated by nonlinear regression analysis using GraphPad Prism (version 9.0; GraphPad Software, Inc.).

### 2.6. Cell toxicity assay

Cells (50,000 per well in 96-well plate) were treated with EGCG or DMSO in DMEM and incubated at 37°C in 5% CO_2_ for 2h. To mimic the conditions used in the infectivity and pseudoparticle assays, media was removed, cells were washed with PBS, and supplemented with normal culture media of DMEM with 10% FBS, 50 U/mL penicillin, and 50 μg/mL streptomycin. After 72 h, cell viability was assessed using alamarBlue cell viability reagent (Invitrogen DAL1025) on a fluorescence plate reader (SpectraMax ID3 Multimode Plate reader).

### 2.7. Time-of-addition experiments

Near-confluent Huh7 and A549 cell monolayers in 12 or 24-well plates were pre-treated with EGCG or DMSO vehicle for 1 h at 37°C in DMEM. Cells were washed with DMEM and infected with 50-200 PFU of HCoV-229E or HCoV-OC43 for 2 h at 33°C. Alternatively, Huh7 and A549 cells were first infected with HCoV-229E or HCoV-OC43 for 2 h at 33°C. After removing the inocula, the infected cells were overlaid with 1.2% microcrystalline colloidal cellulose in DMEM-2% FBS containing EGCG or DMSO. Infectivity was assessed by plaquing efficiency or immunostaining.

### 2.8. Binding assays

OC43 (4 × 10^4^ PFU; MOI 0.05) or VSV-SARS-CoV-2 (2 × 10^5^ PFU; MOI 0.5, chosen to match
 the conditions of the infectivity assay) were exposed for 10 min at 37°C to EGCG or DMSO in DMEM and adsorbed onto pre-chilled Huh7 and A549 cells for 1 h at 4°C. After three washes with cold phosphate-buffered saline (PBS), cells were lysed and RNA was isolated using the Monarch Total RNA Miniprep Kit (NEB T2010S), according to the manufacturer’s instructions. The RNA was transcribed to cDNA using the High Capacity cDNA Reverse Transcription Kit (Applied Biosystems LS4368814). Quantitative real-time PCR was performed on the QuantStudio 3 (Applied Biosystems) instrument, using PowerTrack SYBR Green Master Mix (Applied Biosystems LSA46110), with primers to detect actin (F: 5’-CTGGGAGTGGGTGGAGGC-3’, R: 5’-TCAACTGGTCTCAAGTCAGTG-3’) and OC43 N gene (F: 5’-CCCAAGCAAACTGCTACCTCTCAG-3’, R: 5’-GTAGACTCCGTCAATATCGGTGCC-3’) or VSV N gene (F: 5’-GATAGTACCGGAGGATTGACGACTA-3’, R: 5’-TCAAACCATCCGAGCCATTC-3’). Following normalization to actin, the percentage of binding was expressed relative to binding of virions treated with the DMSO control.

## 3. Results

### 3.1. EGCG inhibits murine and human coronavirus infection

We first tested the effect of EGCG on the infectivity of human common cold CoVs HCoV-229E (an alphacoronavirus) and HCoV-OC43 (a betacoronavirus). HCoV-229E and HCoV-OC43 virions were treated with DMSO or varying concentrations of EGCG for 10 min at 37°C and used to inoculate infection-susceptible human hepatoma Huh7 or human lung epithelial A549 cell monolayers. After 2 h of infection, inocula were removed and cells were overlaid with plaquing media. Infectivity was assessed by plaquing efficiency on Huh7 for HCoV-229E or by immunofluorescence focus-forming assay on Huh7 and A549 cells for HCoV-OC43. EGCG inhibited infectivity of HCoV-229E (IC_50_ = 0.77 μM) and HCoV-OC43 (IC_50_ = 0.49-0.56 μM) at sub- to low micromolar concentrations (**Figure 1A**), similar to other viruses highly susceptible to inhibition by EGCG, such as HSV-1 (Colpitts and Schang, 2014). We next tested the inhibitory activity of EGCG against a murine CoV, murine hepatitis virus (MHV-A59), on murine fibroblast L929 cell monolayers. EGCG inhibited MHV-A59 infection with similar potency as for the human common cold CoVs (IC_50_ = 0.71 μM) (**Figure 1B**) and inhibited the formation of syncytia (**Figure 1C**). Consistent with previous reports, we found EGCG to have minimal effect on cell viability at the low micromolar range in which antiviral activity is exerted (**Figure 1D**).

**Figure 1.**
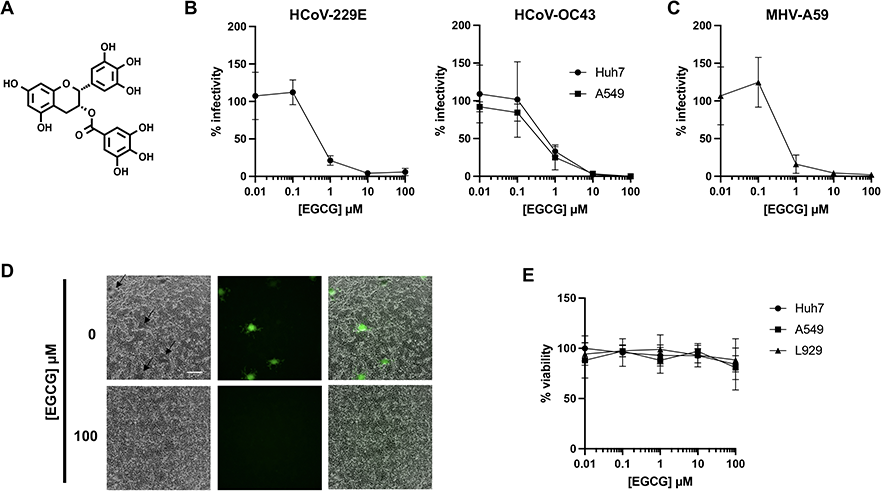
EGCG pre-treatment inhibits infection by diverse CoVs. **(A)** Structure of epigallocatechin gallate (EGCG). **(B-C)** Pre-treatment of authentic HCoV-229E, HCoV-OC43 or murine coronavirus MHV-A59 virions with EGCG for 10 min at 37°C inhibits infection of Huh7, A549 or L929 cells, respectively. Mean values with standard deviation of three independent experiments are plotted. **(C)** A reduction in syncytia (black arrows) was observed for EGCG-treated MHV-A59. Scale bar: 200 μm. **(D)** Cell exposure to EGCG during the 2 h infection had minimal effect on cell viability. Mean values with standard deviation of two independent experiments with duplicates are plotted.

### 3.2. EGCG inhibits entry of highly pathogenic and emerging coronaviruses

We produced luciferase-encoding lentiviral particles pseudotyped with SARS-CoV-1, SARS-CoV-2 and MERS-CoV spike (S) proteins to investigate the effect of EGCG on entry of highly pathogenic CoVs. We also generated lentiviral particles pseudotyped with the spike protein of WIV1-CoV, a bat CoV that binds to human ACE2 (Ge et al., 2013). Huh7 cells were incubated with CoV S pseudoparticles pre-exposed to EGCG or DMSO for 10 min at 37°C. Inocula were removed after 2 h and replaced with fresh DMEM. After 72 h, cells were lysed and luminescence was measured. EGCG inhibited SARS-CoV-1, SARS-CoV-2, MERS-CoV and WIV1-CoV pseudoparticle entry with IC_50_ of 2.5, 48.6, 13.7 and 3.8 μM, respectively (**Figure 2A**). We also generated A549 human lung alveolar epithelial cells stably expressing human ACE2, the receptor for SARS-CoV-1 and SARS-CoV-2, and isolated a clone with high levels of ACE2 expression and enhanced susceptibility to SARS-CoV-2 S pseudoparticle infection (**Supplemental Figure 1**). EGCG similarly inhibited entry of SARS-CoV-1, SARS-CoV-2, and WIV1-CoV pseudoparticles into A549-ACE2 B9 cells, with IC_50_ of 15.9, 24.0 and 24.5 μM, respectively (**Figure 2A**).

**Figure 2.**
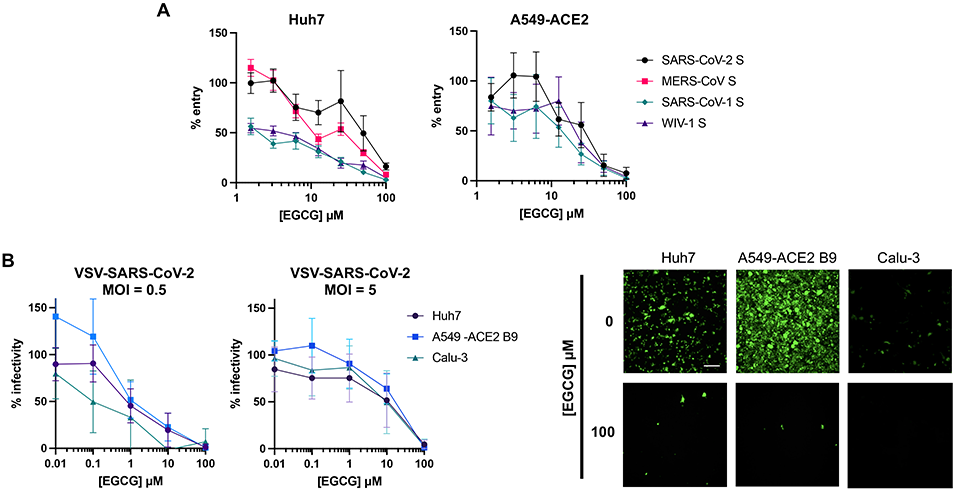
EGCG inhibits entry of highly pathogenic CoVs and an emerging bat CoV. **(A)** EGCG treatment reduced entry of lentiviral particles pseudotyped with S from SARS-CoV-1, MERS-CoV, SARS-CoV-2, or WIV1-CoV. Mean values with standard error of the mean of three independent experiments each done in triplicates are plotted. **(B)** EGCG displayed MOI-dependent inhibition of VSV-SARS-CoV-2 in Huh7, A549-ACE2 B9 and Calu-3. Mean values with standard deviation of three independent experiments each in duplicates are plotted. Representative images are shown. Scale bar: 200 μm.

We assessed the effect of EGCG on replication-competent vesicular stomatitis virus (VSV) virions expressing GFP and SARS-CoV-2 S instead of the VSV glycoprotein (VSV-SARS-CoV-2) (Case et al., 2020). Huh7, A549-ACE2, and lung epithelial Calu-3 cells were infected with EGCG-treated VSV-SARS-CoV-2 virions (MOI of 5). Infected cells were imaged by fluorescence microscopy after 24 h. Similar to the pseudoparticle assays, EGCG inhibited VSV-SARS-CoV-2 S infection with IC_50_ ranging from 11.45 to 15.40 μM, depending on the cell type (Figure 2B), with minimal cytotoxicity up to 100 μM (**Supplemental Figure 2**). We next evaluated whether the effect of EGCG was dependent on MOI. Using the same assay, but with MOI of 0.5, we found that EGCG inhibited VSV-SARS-CoV-2 infection with IC_50_ ranging from 0.14 to 1.13 μM. These findings demonstrate that the inhibitory activity of EGCG depends on the MOI (**Figure 2B**), consistent with the virions as the proposed target of EGCG (Colpitts and Schang, 2014). This also provides a potential explanation for the difference in IC_50_ for authentic viruses investigated at low MOI (<0.001) (**Figure 1A-B**) compared to lentiviral pseudoparticles (**Figure 2A**). Based on p24 ELISA data for lentiviral particles pseudotyped with SARS-CoV-2 spike in our laboratory, we estimate lentiviral titers of approximately 1.7 × 10^6^ transducing units/mL, resulting in an apparent MOI of 3.4 in these experiments.

### 3.3. EGCG acts on virions to inhibit coronavirus binding to cell surfaces

To confirm that EGCG acts on CoV virions, and not on cells, we performed time-of-addition experiments. Cells were pre-treated with EGCG or DMSO 1 h at 37°C, at which point media was removed and cells were infected with HCoV-229E or HCoV-OC43. Alternatively, cells were infected, and EGCG- or DMSO-containing medium was added at 2 h post-infection. Infectivity was determined by plaque assay or immunostaining. In either condition, no inhibition of infectivity was observed until the highest concentration of 100 μM EGCG (**Figure 3A**), consistent with the main effect of EGCG being on viral entry, with virions as the target. To test if EGCG inhibits CoV attachment, we performed binding assays in which cells were incubated with pre-treated virions at 4°C for 1 h to allow for attachment, but not fusion or entry. After cell washing, bound virions were quantified by RT-qPCR (**Supplemental Figure 3**). Pre-chilled Huh7 or Calu-3 cells were inoculated with HCoV-OC43 or VSV-SARS-CoV-2 treated with DMSO, 10 μM EGCG, or 20μg/mL heparin, as a control attachment inhibitor (Clausen et al., 2020). As expected, heparin partially blocked VSV-SARS-CoV-2 binding. We also observed that heparin reduced HCoV-OC43 binding, which unlike other coronavirus does not have a known protein receptor but depends on sialic acid, and, as recently reported, glycosaminoglycans for entry (Wang et al., 2021). EGCG similarly reduced HCoV-OC43 and VSV-SARS-CoV-2 binding to Huh7 and Calu-3 cells (**Figure 3B-C**).

**Figure 3.**
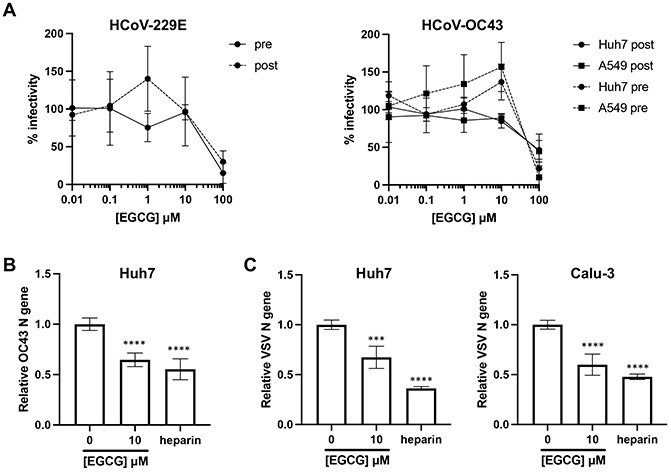
EGCG inhibits CoV attachment to cells. **(A)** Pre-treatment of cells, or treatment of cells post-infection with HCoV-229E or HCoV-OC43, does not inhibit infection to similar extents as virion pre-treatment. Mean values with standard deviation of three independent experiments are plotted. **(B)** HCoV-OC43 virions were pre-treated with EGCG or heparin for 10 min at 37°C then cooled on ice and added to pre-chilled Huh7 for 1 hour on ice. **(C)** VSV-SARS-CoV-2 were pre-treated as described above and added to pre-chilled Huh7 or Calu-3 cells. Attached virions were quantified by RT-qPCR after washing cells with PBS three times. Mean values with standard deviation from qPCR triplicates of two independent experiments are plotted.

## 4. Discussion

Green tea catechins, especially EGCG, exert antiviral effects against diverse viruses. We show that EGCG broadly inhibits CoV infections, and that its activity is due, at least in part, to blocking CoV attachment to target cells. Infection by authentic murine and human CoVs, as well as entry of lentivirus particles pseudotyped with spike proteins of highly pathogenic or potentially emerging CoVs (Ge et al., 2013), was inhibited by EGCG treatment. Interestingly, EGCG was recently shown to significantly inhibit authentic SARS-CoV-2 infection, and time-of-addition experiments suggested that EGCG at least partially blocks SARS-CoV-2 entry (Henss et al., 2021), consistent with our findings. Here, we expand the scope of EGCG activity to other CoVs, including murine and bat CoVs, and demonstrate that EGCG inhibits CoV attachment and infection of lung epithelial cell lines.

A variety of mechanisms for the antiviral activity of EGCG have been described. EGCG inhibits the primary attachment of diverse viruses including VSV, IAV, HCV, HSV-1 and vaccinia virus (Colpitts and Schang, 2014). EGCG is proposed to act directly on HCV virions to block attachment in a genotype independent manner, with no alteration in HCV (co)-receptor expression nor any effect on viral replication, assembly, or release (Calland et al., 2012; Ciesek et al., 2011). For HBV, EGCG may inhibit entry (Huang et al., 2014), as well as replication, with proposed roles in impairing HBV replicative intermediates during DNA synthesis (He et al., 2011), diminishing the transcriptional activation of the HBV core promoter (Xu et al., 2016), and enhancing lysosomal acidification to adversely affect HBV replication (Zhong et al., 2015). EGCG is suggested to prevent the binding of HIV glycoprotein 120 to its receptor molecule CD4 on cells (Williamson et al., 2006), and to act as an HIV reverse transcriptase inhibitor (Li et al., 2011). Finally, recent studies have identified *in vitro* inhibitory effects of EGCG against the SARS-CoV-2 3C-like protease (Chiou et al., 2021; Du et al., 2021; Jang et al., 2021, 2020). However, our time-of-addition experiments with HCoV-229E and HCoV-OC43, as well as those of others with SARS-CoV-2 (Henss et al., 2021), strongly suggest that EGCG predominantly inhibits authentic CoV infection by blocking entry, although other antiviral activities might be exerted at higher concentrations. Furthermore, we show that EGCG reduces cell surface binding of HCoV-OC43 and VSV-SARS-CoV-2, in a similar manner to heparin, which has been demonstrated to competitively block virion attachment (Clausen et al., 2020).

While COVID-19 vaccine development and deployment are ongoing, extensive efforts are underway to identify antibodies, peptides, and small molecules to prevent SARS-CoV-2 entry (Xiu et al., 2020). To protect against emerging CoVs, the identification of broad-spectrum entry inhibitors is a priority. Recently, lactoferrins have been described to disrupt CoV primary attachment, mediated by heparan sulfate interactions, conferring antiviral activity against multiple CoVs *in vitro*, demonstrating the potential for a pan-coronavirus inhibitor (Hu et al., 2021). Small molecules continue to be preferred as drugs in general due to superior pharmacokinetic properties and simpler synthesis (Ngo and Garneau-Tsodikova, 2018). As with other green tea catechins, EGCG does not accumulate at high levels, is unstable under physiological conditions, and is rapidly metabolized (Chow et al., 2003; Lambert et al., 2003; Smith, 2011). However, the observation that a small molecule exerts pan-coronavirus activity provides the basis for the development of entry inhibitors that could provide an effective antiviral strategy to protect against future emerging CoV infections.

## Acknowledgements

The following reagents were obtained through BEI Resources (NIAID, NIH): *Homo sapiens* A549 Lung Carcinoma Cells (NR-52268); Murine 17CL1 Cell Line (NR-53719); SARS-Related Coronavirus 2, Wuhan-Hu-1 Spike-Pseudotyped Lentiviral Kit (NR-52948); Human Coronavirus OC43 (NR-52725) and 229E (NR-52726). We thank Gerard O’Leary for Python processing scripts, Dr. Selena Sagan (McGill University) for providing coronavirus infection protocols, and Dr. Chantelle Capicciotti (Queen’s University) for helpful discussions.

