## Supplementary Figures for "The green tea catechin EGCG provides proof-of-concept for a pan-coronavirus entry inhibitor"

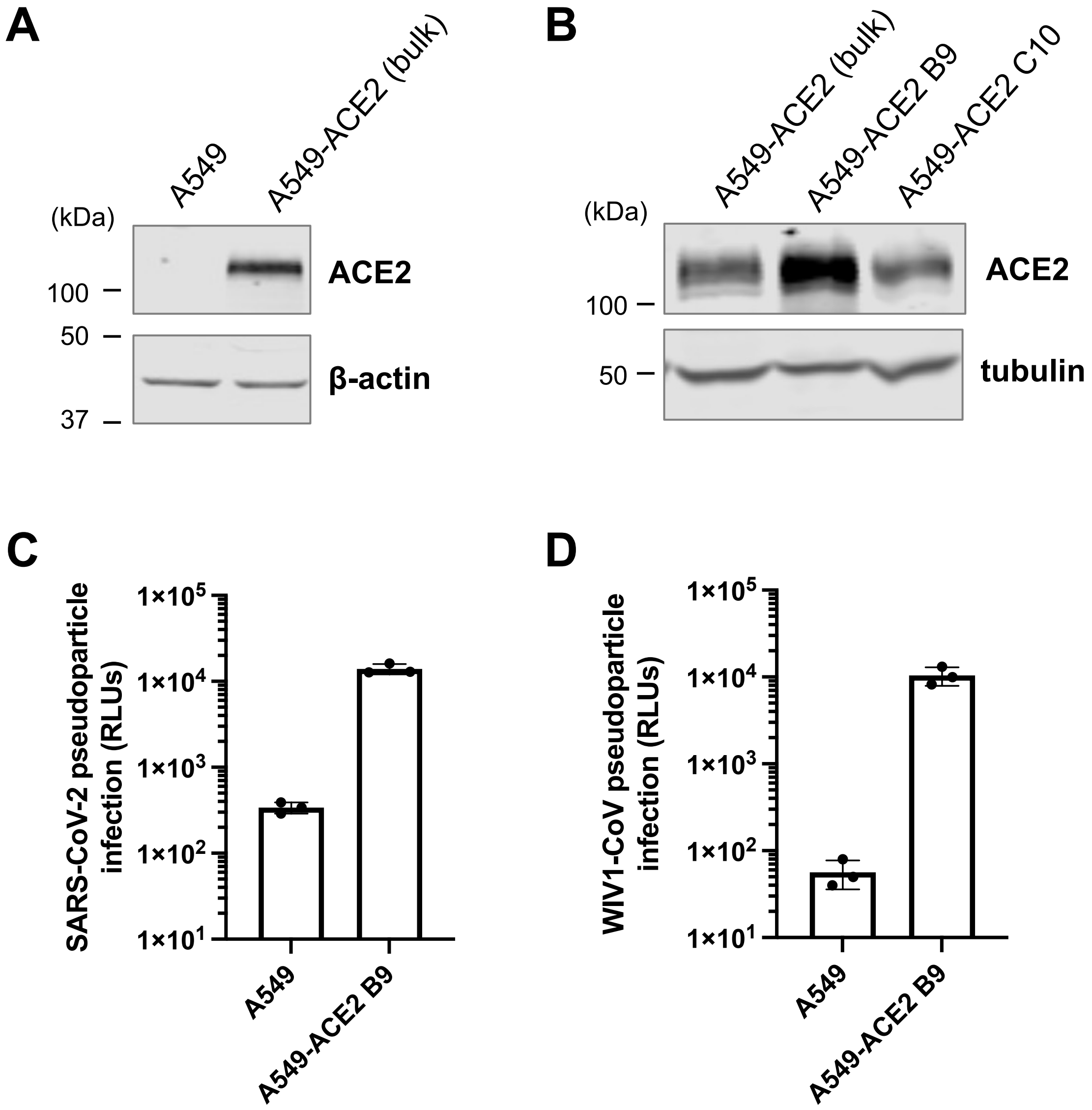


**Figure S1. A549-ACE2 B9 expresses high levels of ACE2 and is susceptible to SARS-CoV-2 pseudoparticle infection.** Western blot shows overexpression of ACE2 in the bulk transduced A549 cells **(A)** and in single cell clones **(B)** with A549-ACE2 B9 showing highest levels of ACE2 expression. Representative blots are shown. SARS-CoV-2 **(C)** and WIV1-CoV **(D)** pseudoparticle infection is dependent on ACE2. Mean value with standard deviation are plotted.

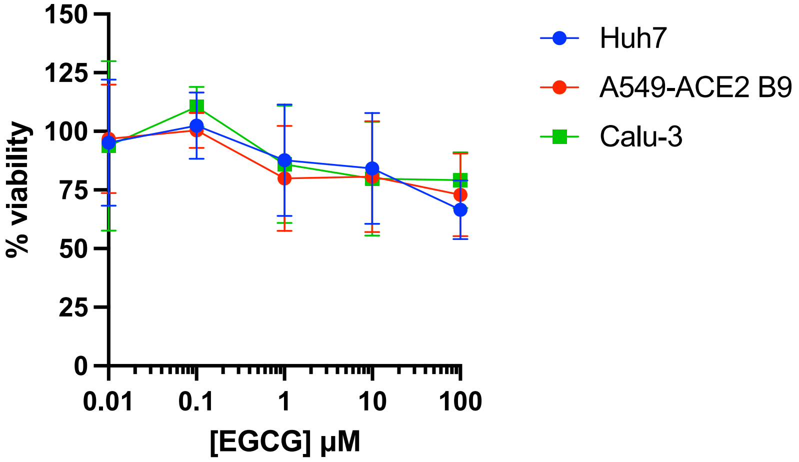


**Figure S2. EGCG exposure for 24 h has minor effects on cell viability.** Cell viability was assessed after 24 h EGCG exposure. Mean values with standard deviation of two independent experiments with duplicates are plotted.

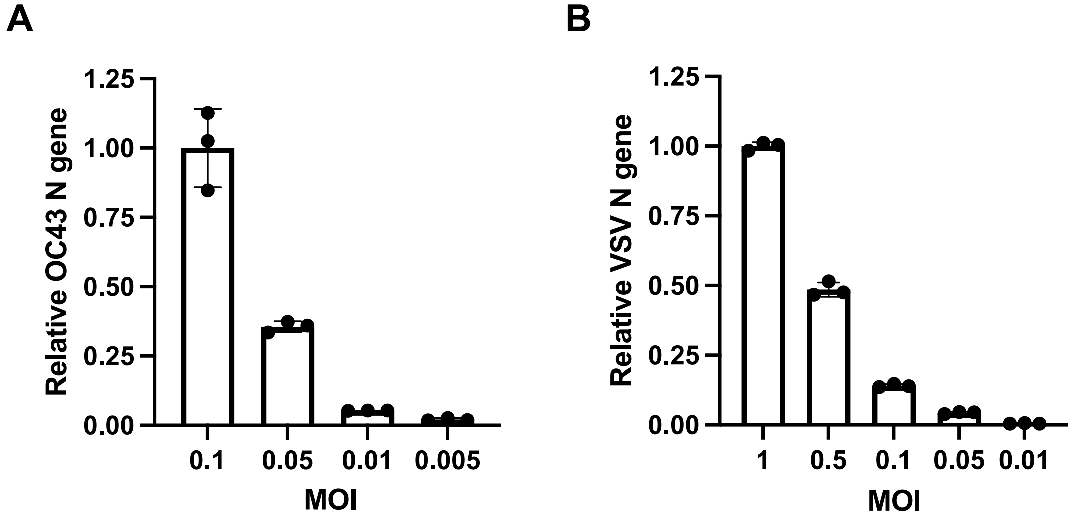


**Figure S3. RT-qPCR quantification of bound virions.** Pre-chilled Huh7 cells were inoculated with OC43 **(A)** or VSV-SARS-CoV-2 (B) at indicated MOI for 1 hour on ice. Attached virus was quantified by RT-qPCR after washing with PBS three times. Mean values with standard deviation from qPCR triplicates are plotted.
